# Experimental evolution of extremophile levels of radiation resistance in *Escherichia coli*

**DOI:** 10.1101/2021.10.18.464883

**Authors:** Steven T. Bruckbauer, Benjamin B. Minkoff, Takeshi Shinohara, Anna Lipzen, Jie Guo, Elizabeth A. Wood, Michael R. Sussman, Christa Pennacchio, Michael M. Cox

## Abstract

Recent human development of high-level sources of ionizing radiation (IR) prompts a corresponding need to understand the effects of IR on living systems. One approach has focused on the capacity of some organisms to survive astonishing levels of IR exposure. Using experimental evolution, we have generated populations of *Escherichia coli* with IR resistance comparable to the extremophile *Deinococcus radiodurans*. Every aspect of cell physiology is affected. Cellular isolates exhibit approximately 1,000 base pair changes plus major genomic and proteomic alterations. The IR resistance phenotype is stable without selection for at least 100 generations. Defined and probable contributions include alterations in cellular systems involved in DNA repair, amelioration of reactive oxygen species, Fe metabolism and repair of iron-sulfur centers, DNA packaging, and intermediary metabolism. A path to new mechanistic discoveries, exemplified by an exploration of *rssB* function, is evident. Most important, there is no single molecular mechanism underlying extreme IR resistance.

## Introduction

Ionizing radiation (IR), particularly from gamma rays, induces severe damage to all cellular macromolecules. Gamma radiation works primarily via H_2_O radiolysis to produce reactive oxygen species (ROS) (*1, 2*). Direct ionization and indirect oxidation of DNA through ROS leads to the accumulation of double-strand breaks (DSBs) of genomic DNA, which are lethal if left unrepaired (*3–7*). Cellular proteomes and lipidomes are also damaged (*8–16*). Some organisms exhibit an extraordinary capacity to survive high doses of IR. Of seminal interest has been the bacterium *Deinococcus radiodurans*, which was first isolated due to its ability to survive IR-based sterilization of canned meat (*17*). Whereas a dose of 3-5 Gy is lethal to a human, *Deinococcus* can survive 5,000 Gy without lethality (*18–21*). Research has revealed several mechanisms utilized by *D. radiodurans* to survive high levels of IR exposure, including specialized mechanisms of DNA end protection and DSB repair, and a notable ability to accumulate Mn^2+^ions as a means of ameliorating the ROS generated by IR (*22–30*).

For most of the history of our planet, there have been no environments that would expose living organisms to high levels of ionizing radiation. That situation has changed with the advent of nuclear power, X-rays, the prospect of extended spaceflight, and more. Understanding how cells respond to ionizing radiation becomes more important as the potential for IR exposure increases.

Our understanding of IR resistance is rudimentary at best. The study of extremophiles such as *D. radiodurans* has been illuminating but can never provide a complete view of IR resistance mechanisms. The reason is simple. The evolution of *Deinococcus* was not driven by ionizing radiation. Prior to its discovery in the 1950s (*17*), this organism was never exposed to the requisite IR-laden environments (*31*). Instead, *Deinococcus* is a desert dweller and evolved to survive desiccation (*32–35*). IR resistance is merely a byproduct of that parched origin. It is thus unlikely that evolution has equipped *Deinococcus* with all possible biological strategies for IR resistance. It is not clear that we know what to look for with respect to additional possible mechanisms. It is not clear that we even understand all the strategies embodied in Deinococcus. More broadly, no research has ever tested the limits. How resistant to ionizing radiation can an organism become?

An alternative and unbiased approach to defining mechanisms of IR resistance, pioneered for this phenotype by Evelyn Witkin in 1946 (*36, 37*), is experimental evolution. Take a species that is not IR resistant, convert it into something that is IR resistant, and see what changes. We are not the first to take this approach (*38–40*). However, previous work was carried out before the development of “omics” technologies that could efficiently identify cellular changes associated with the phenotype.

With a goal of elucidating new mechanisms of IR resistance, we are pursuing a long-term evolution experiment with the goal of generating IR resistance in four populations of *E. coli* equivalent to and eventually surpassing *D. radiodurans*. After 150 cycles of selection carried out over 5 years, our initial goal of IR resistance parity with *Deinococcus* has been achieved. Earlier reports of results after 50 or 100 cycles of selection have described important intermediate steps in this project (*41, 42*). Here, we present *E. coli* strains with documented extremophile levels of IR resistance. Two of our four replicate populations of evolved *E. coli* now match the survival of *D. radiodurans* out to a dose of 6,000 Gy of IR, whereas doses in excess of 1,000 Gy effectively sterilize cultures of the Founder *E. coli* strain. Combined genomic and proteomic approaches have revealed that changes to systems involved in DNA repair, ROS amelioration, Fe metabolism, and aerobic metabolism have partially driven acquired IR resistance in at least one of the four evolved populations. A path to describing additional mechanisms is evident. In addition, we demonstrate that an elevated stress response contributes to IR resistance. The overall effort highlights the lack of a single molecular mechanism underlying biological resistance to ionizing radiation. Layers of contributing mechanisms are evident within these four evolved *E. coli* populations.

## Results and Discussion

### Ionizing radiation resistance has matched that of Deinococcus radiodurans

To summarize our experimental evolution protocol (detailed in the *Materials and Methods*), at each cycle of selection, each of the four replicate populations is initially grown to early exponential phase and cultures are washed with PBS to remove all nutrients or potential ROS-ameliorating agents present in growth media. The washed cultures are treated with sufficient IR to kill 99% of each of the four replicate populations at each cycle of selection (thus the dose used increases as IR resistance increases). The irradiation utilizes a clinical linear accelerator (Linac) dosing at 70 Gy/min. Following irradiation, a portion of each culture is used to determine percent survival and the rest is allowed to recover in fresh growth medium overnight before storage of the new population at –80 °C the following day. This protocol has resulted in four evolved lineages (IR9, IR10, IR11, and IR12), with distinct populations at each cycle of selection (i.e. IR9-150 is the population from the IR9 lineage at round 150 of selection), and 10 isolates from each population (i.e. IR9-150-1 and IR9-150-2 are two separate isolates from population IR9-150). We note that population IR9 has two sub-populations which have competed in clonal interference for about 80 selection cycles through round 150 (*42*). Isolates IR9-150-1 and IR9-150-2 are taken from different sub-populations and exhibit many genetic and proteomic distinctions in spite of their presence in the same population.

After beginning with a dose of 500 – 750 Gy, at round 150 of selection a dose of 3700-4000 Gy is required to achieve 99% killing **(Figure S1)**. Furthermore, two populations, IR9-150 and IR10-150, now exhibit survival curves equivalent to *D. radiodurans*, where cultures of each population and *D. radiodurans* have approximately 0.1% survival at a dose of 6000 Gy **(Figure 1)**.

**Figure 1.**
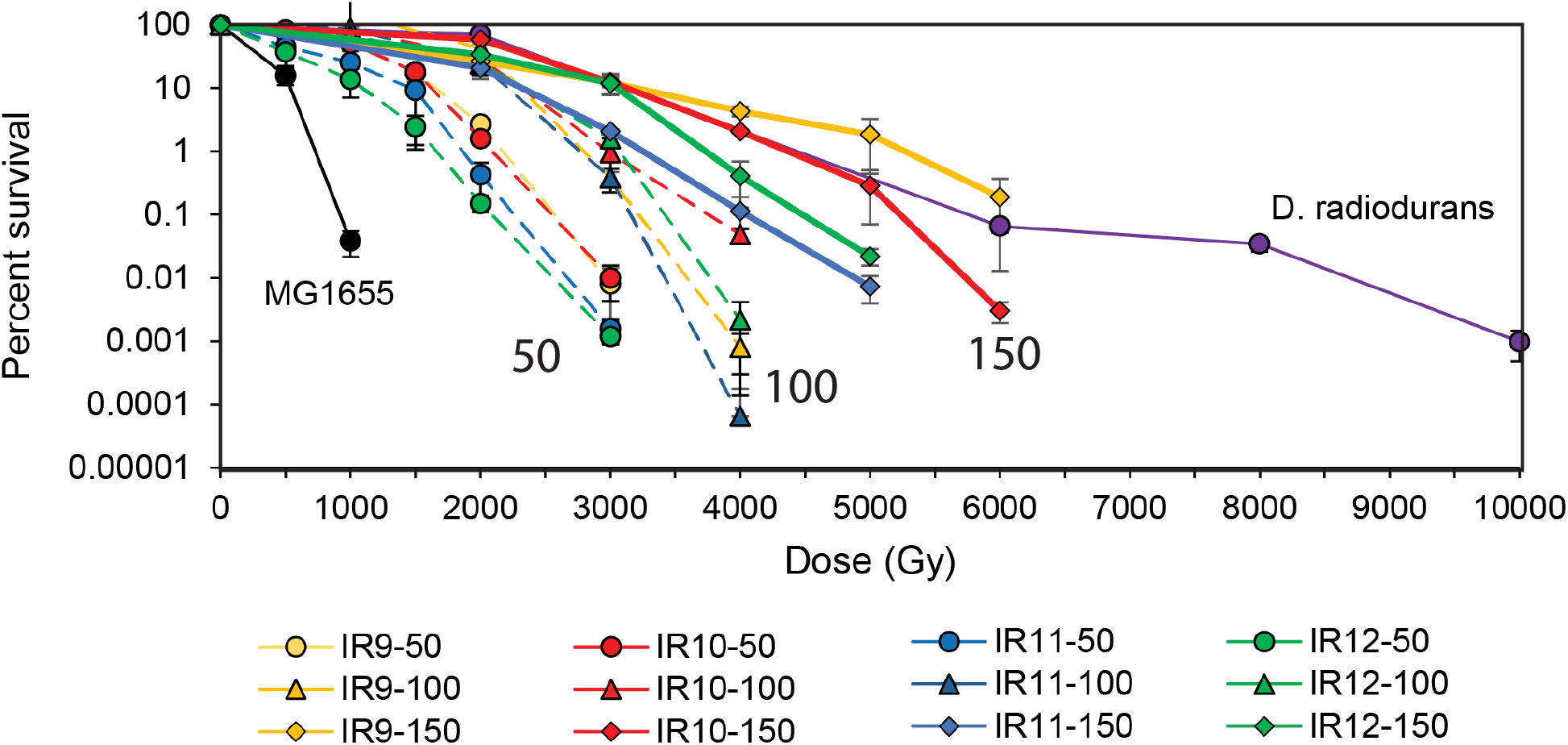
After 150 cycles of selection experimentally evolved *E. coli* populations exhibit IR resistance comparable to *D. radiodurans*. Survival curves of evolved populations compared to previously evolved *Escherichia coli* populations and *Deinococcus radiodurans*. MG1655 is the Founder strain used to begin the evolution experiment. IR’X’-50 and IR’X’-100 are populations of the indicated lineages after 50 or 100 cycles of selection with IR. Early exponential phase cultures of the indicated strains were exposed to electron beam IR as described in the *Materials and methods*. Error bars represent the standard deviation of CFU/mL calculations from a single experiment performed in biological triplicate. Survival data for MG1655, populations after 50 cycles of selection, and *D. radiodurans* strain R1 at 2000 and 6000 Gy are as previously reported (*41, 42*).

Radioresistance in these four populations is a stable phenotype. After growth for ~100 generations in the absence of IR exposure, isolates from each of these populations maintains the same level of IR resistance (**Figure S2)**. However, these isolates are not free of fitness tradeoffs. Each isolate has clear growth rate defects **(Figure S3)** as well as readily observable changes in cell morphology **(Figure S4)**. Such differences are expected, as after 150 cycles of selection isolates contain about 1000 single-nucleotide polymorphisms (SNPs) and other small changes on average **(Table S1)**. Larger genomic alterations are evident in the populations as well. Some were reported previously (*42*). A new genomic deletion present in IR11 is added in **Figure S5**.

### Evolved isolates exhibit enhanced DNA repair and resistance to protein oxidation

DNA double strand breaks accumulate with increasing doses of IR and must be repaired in order for the exposed cell to survive. Thus, we assayed IR-mediated damage to genomic DNA and the dynamics of DNA repair in evolved isolates from each population after a dose of 1000 or 4000 Gy, utilizing Pulsed-Field Gel Electrophoresis (PFGE). All evolved isolates demonstrated a clear capacity to repair shattered genomic DNA at a dose of 4000 Gy, attesting to enhanced capacity for DNA repair. In contrast, the Founder strain did not recover from this dose **(Figure 2)**. These results align with our previous observations that by rounds 50 and 100 of selection, numerous mutations in DNA repair proteins (RecA, RecD, RecN, and RecJ) were drivers of evolved IR resistance (*41, 42*). Many of these mutants affect proteins involved in double strand break repair. The evolved isolates exhibited different rates of DNA repair. Repaired genomic DNA is apparent in isolate IR12-150-1 only 3 hr post-irradiation with 4000 Gy. Furthermore, IR11-150-1 and IR12-150-1 appear to better protect their genomic DNA from IR-mediated DSBs, where cells harvested immediately post-irradiation with 1000 Gy contain at least some more or less intact genomes. Thus, it appears that the evolved lineages may have developed differing mechanisms of maintaining genome stability despite equivalent IR exposure.

**Figure 2.**
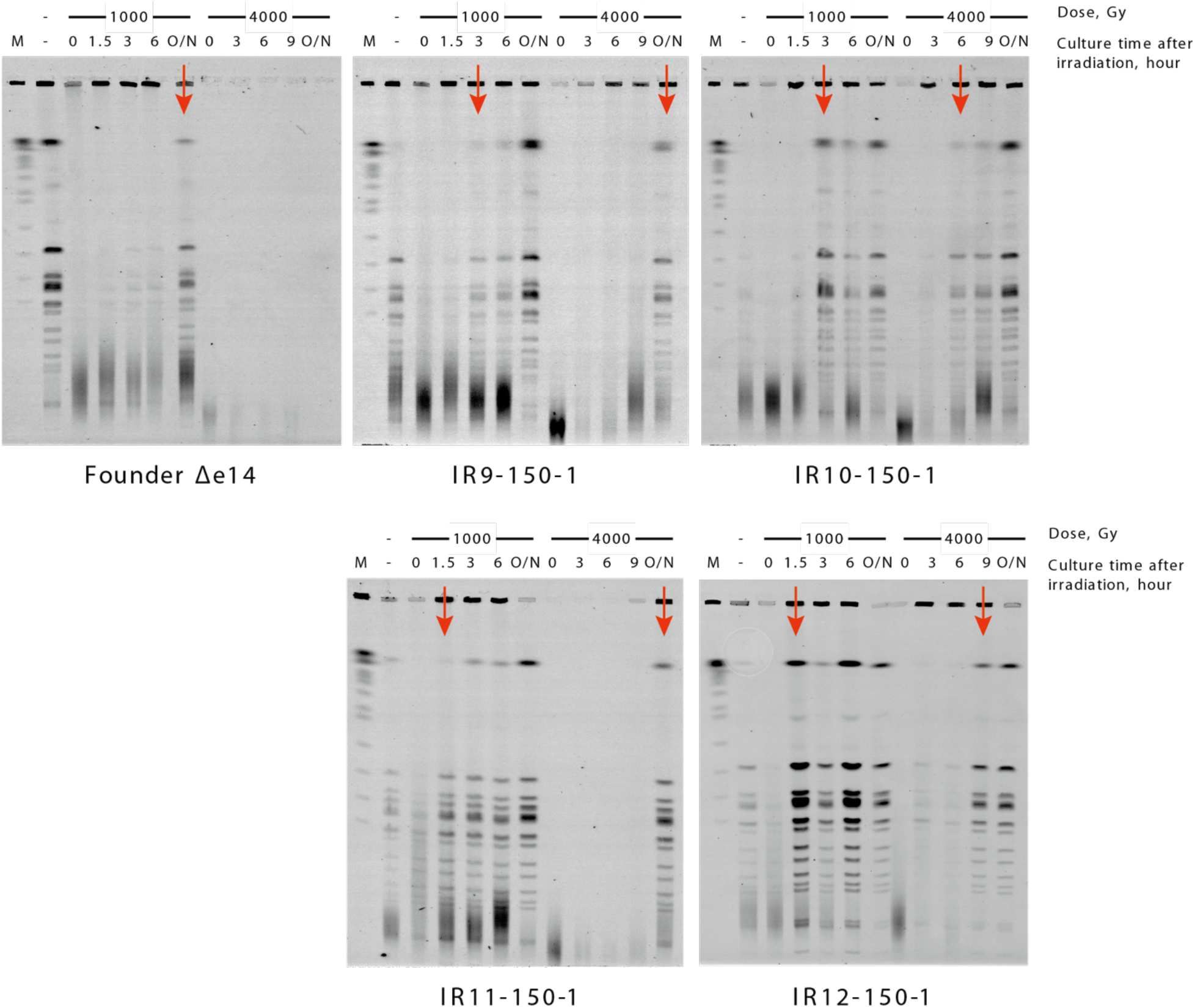
Evolved isolates experience and repair extensive IR-induced DNA damage. Pulsed-field gel electrophoresis (PFGE) was used to assay the extent and repair of DNA damage induced by IR exposure at 1000 and 4000 Gy. Genomic DNA is digested with restriction enzyme NotI to facilitate observation of intact versus degraded genomes. At 4000 Gy, cultures of MG1655 are completely killed and no repair of genomic DNA is observed. Red arrows indicate the first appearance of a banding pattern suggesting intact genome. Results are representative of two independent experiments. PFGE was performed as described previously (*61*), and as in *Materials and methods*.

### Proteomics and genomics provide a new entrée to mechanisms of evolved IR resistance

The four evolved populations were subjected to deep sequencing to identify all mutations present in more than 2% of the cells **(Supplementary Dataset S1).** Genes subject to mutation in multiple populations are of particular interest as potential candidates for phenotype contributions, and these are listed in **Table S2**. In parallel, using mass spectrometry approaches we have described previously (*10, 11*), we surveyed the proteomes of the Founder wild type strain and two evolved isolates from the current experiment. Analysis of the quantified IR9-150-1 and IR9-150-2 proteomes at early exponential phase growth under normal growth conditions (no IR) revealed significant changes to the composition of their proteomes compared to MG1655 (**Figure 3**). With a significance threshold of at least a 2-fold increase or decrease (adjusted p-value < 0.05), each isolate had over 300 proteins with altered abundance (IR9-150-1: 156 proteins increased and 165 decreased; IR9-150-2: 150 proteins increased and 317 proteins decreased). Such changes potentially reveal what pathways are beneficial or expendable for IR resistance.

**Figure 3.**
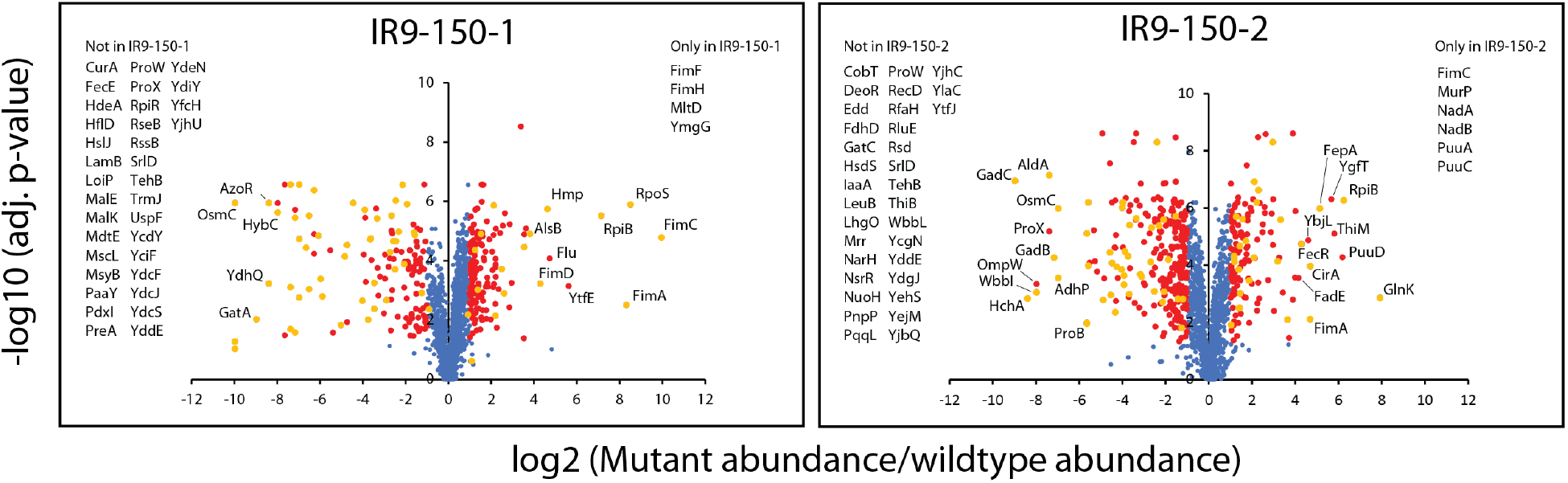
The proteome composition of IR9-150-1 and IR9-150-2 have diverged from *E. coli* MG1655. Volcano plots depict each protein detected with a circle, with the fold difference in abundance compared to the same protein in MG1655 on the x-axis and the p-value of that change on the y-axis. Proteins with a significant fold change are colored in red. Numerous proteins were not detected, or only detected in the evolved isolates. Such proteins are listed next to their respective volcano plots, as they cannot be plotted due to no actual fold increase or decrease. The proteome composition of MG1655, IR9-150-1, and IR9-150-2 was determined utilizing label free quantification mass spectrometry (LFQ-MS). Significant changes in protein abundance were defined as an increase or decrease greater than two-fold, with adjusted p-values less than 0.05 (calculated using Benjamini-Hochberg correction). Supplementaary Datasets 1 and 2 contain the genomics and proteomics data used for these analyses, respectively.

Across both isolates, large changes in abundance of greater than 10-fold (observed in 18 proteins increasing and 52 proteins decreasing in IR9-150-1, and 24 proteins increasing and 42 decreasing in IR9-150-2) are largely explained through genomic sequencing data. Large increases in abundance are often due to mutations in their cognate regulatory proteins. For example, the pentose phosphate pathway protein RpiB is increased due to a frameshift in its regulator, RpiR (also called AlsR) (*43, 44*); pilus proteins from the *fim* operon are likely increased due to mutations in the FimE site-specific recombinase, locking the *fim* promoter in the ‘on’ orientation (*45–47*); and RpoS is increased in IR9-150-1 due to an early frameshift in its regulator, RssB, that effectively eliminates RssB function (*48–50*). Some increases in protein abundance may be linked to intergenic mutations affecting transcription or translation (such as two mutations at –150 and +21 relative to the transcription start site of *uvrB* in both IR9-150 isolates), or due to regulatory changes (i.e. Fe-starvation, explained below). Large decreases in protein abundance are typically due to introduced stop codons or frameshift mutations, or deletion of the gene encoding the protein in question (particularly due to a > 100 kb deletion in IR9 (*42*)).

In prior work characterizing the evolved lineages after 50 (*41*) and 100 (*42*) cycles of selection, we found that evolved IR resistance appeared to proceed through at least two distinct phases: (1) adaptation of DNA repair pathways in all four lineages, followed by (2) lineage-specific adaptations, potentially focusing at least in part on adaptations to respiration, Fe-S cluster repair, and polyamine metabolism in lineage IR9 (*42*). The proteomics and additional genomic sequencing conducted now at round 150 of selection largely confirm the trends first discovered through genomics datasets, as well as highlight new potential players in evolved radioresistance. In the following analysis, we highlight mutations confirmed to be contributors to IR resistance as well as proteomic changes that suggest additional mechanisms to guide additional investigation.

#### DNA repair

The proteomics effort highlights adaptations of the DNA repair systems that were not documented earlier. Each of the two round 150 isolates have increased levels of proteins involved in nucleotide excision repair (UvrB, LigA) as well as proteins involved in remodeling DNA (RuvA, GyrA, GyrB, ParC), suggesting new ways to bolster a cells capacity to process damaged DNA substrates **(Figure S6)**. In conjunction with increased levels of GyrA (IR9-150-2) and GyrB (IR9-150-1), a decrease in the gyrase inhibitor SbmC in both isolates further underlines the potential importance of increased gyrase activity in radioresistance. Additional decreases in DNA repair proteins focus on exo and endonucleases (RecD, RecJ, SbcD, Nth). As IR-induced double strand breaks increase, the capacity of exonucleases to wreak genomic havoc also increases. Hence, a reduction in cellular nuclease activity may contribute to the maintenance of genome integrity in a high IR environment. Mutations in proteins involved in double strand break repair (RecD, RecN, RecJ, RecA) make clear contributions to IR resistance, as documented previously (*41*), and the IR resistance conferred by the changes in RecD and RecJ may reflect declines in nuclease function.

#### Response to ROS

ROS generated by IR or respiration have the capacity to not only generate DNA damage, but also oxidize proteins side chains and destroy protein-coordinated Fe-S clusters (*51, 52*). IR9-150-1 and IR9-150-2 exhibit extensive proteomic changes which are suggestive of suppressing ROS formation linked to respiration by reducing the level of associated proteins **(Figure S6)**. Furthermore, both isolates exhibit significant increases in proteins involved in repair of damaged Fe-S clusters, in particular members of the *suf* operon (**Figure S6**). These changes build on previous observations that variants of the ATP synthase component AtpA and Fe-S repair protein SufD were partially responsible for the radioresistant phenotype of lineage IR9 after 100 cycles of selection (*42*).

The new proteomics data show that isolate IR9-150-1 has increased levels of the ROS amelioration enzymes Hmp, SodA, and AhpF in addition to a large increase in the stress response sigma factor RpoS (which in turn controls transcription of the ROS amelioration proteins PqiA, PqiB, PqiC, and KatE). All of these changes have the potential to reduce the oxidative damage to cellular constituents of all kinds. In contrast, numerous ROS-responsive proteins are observed at decreased levels in IR9-150-2 (KatG, PqiB, PqiC, OxyR, Dps) suggesting that the two sub-populations of IR9 are pursuing quite distinct paths to IR resistance.

#### Fe metabolism

In addition to proteomic confirmation of adaptations previously identified through genomics, the proteome of IR9-150-2 exhibits a strong indication of altered Fe metabolism. Pathways focused on the production and uptake of Fe-scavenging siderophore enterobactin are all increased in abundance, while Fe-storage proteins Dps and Bfr are both decreased. The changes to Fe metabolism may be related to a frameshift in the ExbB protein (ExbB S67fs), which would additionally knock ExbD out of frame. The ExbBD protein complex (in conjunction with TonB) provides the mechanical energy necessary to facilitate enterobactin uptake across the outer membrane. Without these proteins cells can no longer utilize enterobactin as a means of Fe acquisition, leading to Fe starvation (*53, 54*). Interestingly, metals analysis through ICP-MS (**Figure S7**) did not reveal a decreased concentration of Fe in IR9-150-2 cells, suggesting that while an inability to uptake enterobactin may trigger the Fe starvation response, the cells are able to acquire Fe by other means. However, the Mn/Fe ratio was somewhat increased in several of the populations, perhaps telegraphing some evolutionary movement towards an ROS amelioration mechanism characterized by Daly and coworkers (*22–30*).

### An altered stress response through RpoS highlights evolutionary divergence of IR resistance mechanisms

The new proteomic and genomic data highlight many potential new contributions to the IR resistance phenotype, but each requires testing. To illustrate how these evolved populations can be exploited for mechanistic discovery, we further explored the frameshift in gene *rssB*, resulting in increased expression of RpoS. The frameshift occurs early in the gene (at codon 3), essentially deleting *rssB* function. This mutation is present in one sub-population of IR9, exemplified by isolate IR9-150-1, but not in the sub-population from which isolate IR9-150-2 is derived or in population IR10. The *rssB* mutation contributes to IR resistance in a context-dependent manner. When a wild type *rssB* gene is restored in IR9-150-1, while retaining all other mutations in this evolved isolate, there is no evident effect on IR resistance at 3,000 Gy. However, at 4,000 Gy, IR resistance is strongly suppressed (**Figure 4**). Thus, the *rssB* mutation is important in this genetic background primarily at high doses of IR. When an *rssB* deletion was engineered into isolate IR9-150-2, no increase in IR resistance was observed. Instead, the change proved to be slightly deleterious (Figure 4). A similar deletion engineered into an isolate from population IR10 (IR-150-1) had no significant effect on IR resistance. An *rssB* deletion introduced into a wild type background produced a significant increase in IR resistance at 1,000 Gy (**Figure S8**), providing additional confirmation that this alteration can contribute to the IR resistance phenotype. Notably, elevated expression of RpoS in a wild type background can slow growth under at least some conditions (*55*), an effect that may have constrained the emergence of similar mutations earlier in the experimental evolution trial. The mutation does not measurably add to the IR resistance of IR9-150-2 or IR10-150-1, indicating that its effects depend on genomic context.

**Figure 4.**
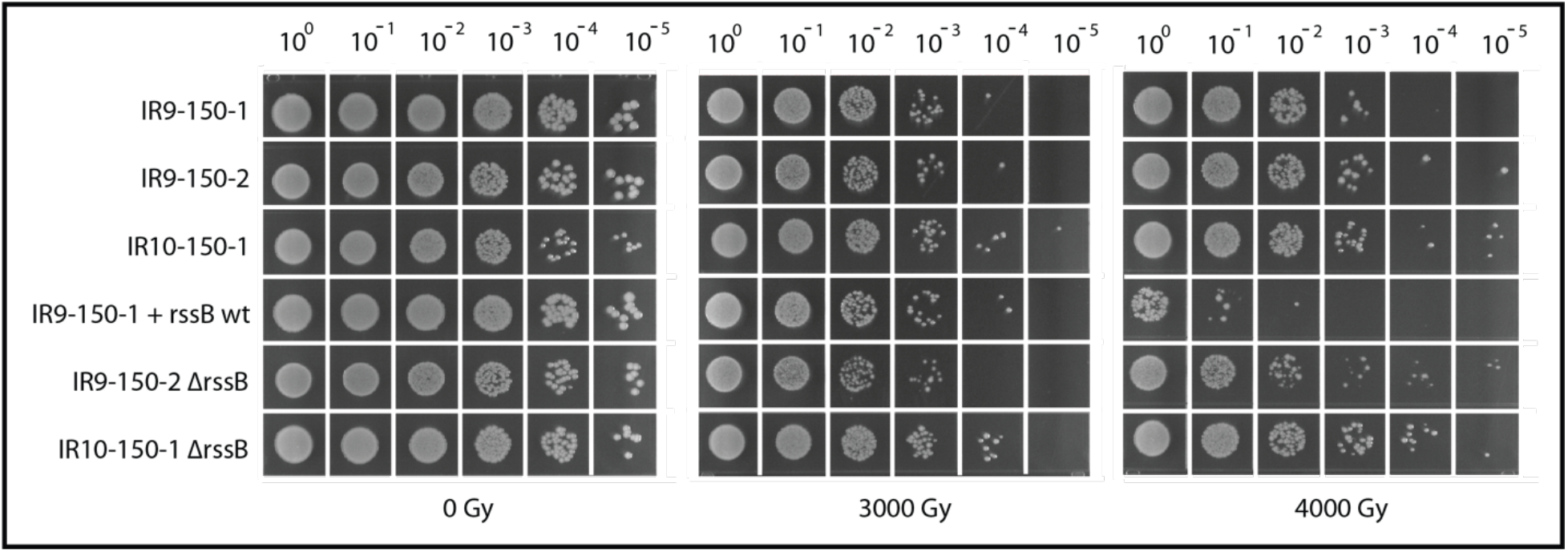
Loss of *rssB* enhances IR resistance in a context-dependent manner. When *rssB* is converted to the wildtype sequence in IR9-150-2, a loss of IR-resistance is observed at 4000 Gy. However, at 3000 Gy no effect is observed from loss of the frameshifted *rssB*, highlighting the dose-dependent context of some beneficial mutations. Furthermore, when *rssB* is deleted from IR9-150-2, or IR10-150-1 we see no effect (IR10-150-1) or deleterious phenotypes (IR9-150-2) at the doses tested. Therefore, beneficial effects from the loss of *rssB* is also dependent on genomic context (see Figure S8 for the effects of an *rssB* deletion in a wild type background.

Whereas we have identified quite a number of mutations that contribute to IR resistance in the present study and previously (*41, 42*), we cannot yet account for the complete phenotype in any one population or sub-population. Some obvious candidates for additional study are revealed here and more will be identified. As exemplified by the *rssB* mutation, these experimentally evolved populations provide a robust platform for the discovery of new mechanisms of IR resistance, although complexity is evident. This study also makes several more general and important points. First, we have not yet reached a dose of ionizing radiation that would preclude survival if appropriate genomic changes are present. The potential for amelioration of IR-mediated damage in living organisms is great and may be greater than that seen in any extant extremophile studied to date. Second, there is not just one way to survive the damage inflicted by IR. Instead, survival may depend on layers of mechanisms that both prevent IR-inflicted damage and facilitate repair of any damage that occurs. The existence of multiple paths to IR resistance may eventually offer multiple points of intervention for cells from bacteria to human that may require this phenotype. Finally, it is likely that at least some of the mechanisms that may contribute to IR resistance involve processes never or rarely discussed in the context of this phenotype. Contributions by alterations in genes involved in DNA packaging (*cadA*) and repair of Fe-S clusters (*sufD*)(*42*), as well as the *rssB* mutation described here, provide examples. Experimental evolution provides an unbiased path to their discovery.

## Acknowledgements

This work was supported by grant GM112757 from the National Institute of General Medical Sciences (NIGMS), and by grants 2817 and 502930 from the Joint Genome Institute, United States Department of Energy. The work conducted by the U.S. Department of Energy Joint Genome Institute, a DOE Office of Science User Facility, is supported by the Office of Science of the U.S. Department of Energy under Contract No. DE-AC02-05CH11231. STB was supported by a Morgridge Biotechnology Scholarship from the Vice Chancellor’s Office for Research and Graduate Education, University of Wisconsin-Madison. BBM was supported by the Department of Defense Defense Threat Reduction Agency grant HDTRA1-16-1-0049.

## Dedication

This paper is dedicated to Evelyn Witkin on the occasion of her 100^th^ birthday.

## Data and Resource Availability Statement

Mutations detected via deep sequencing at round 150 are listed in Supplementary Dataset 1. MS datasets for each strain are available online at the Proteomics Identification Database (https://www.ebi.ac.uk/pride/; accession number: PXD024784) and are included as Supplementary Dataset 2. All data utilized here are incorporated within this publication or in one of these datasets. In addition, all “omics” data, as well as all bacterial populations, isolates, and cellular constructs associated with this project, <u>both published and unpublished</u>, are available to any interested researcher at any time. For a current listing, please inquire.

## Notes

### Competing Interest Statement

The authors have declared no competing interest.

