## Supplemental materials for "Experimental evolution of extremophile levels of radiation resistance in *Escherichia coli*"

### Supplementary Materials

#### Materials and methods

*Growth conditions and bacterial strains used in this study*

Unless otherwise stated, *E. coli* cultures were grown in Luria-Bertani (LB) broth (*56*) at 37°C with aeration. *E. coli* were plated on 1.5% LB agar medium (*56*) and incubated at 37°C. Overnight cultures were grown in a volume of 3 mL for 16 to 18 hr. Exponential phase cultures were routinely diluted 1:100 in 10 mL of LB medium in a 50 mL Erlenmeyer flask and were grown at 37°C with shaking at 200 rpm and were harvested at an $OD_{600}$ of 0.2, unless otherwise noted. After growth to an $OD_{600}$ of 0.2, cultures were placed on ice for 10 min to stop growth before being used for assays.

All *E. coli* strains used for *in vivo* assays in this study are mutants of *E. coli* K-12 derivative MG1655 (*57*). Genetic manipulations to transfer mutations or delete genes were performed as previously described (*58, 59*). *D. radiodurans* is an R1 derivative (ATCC13939) (*17*). Overnight liquid cultures were incubated at 30 °C with shaking for 24 hr; exponential phase cultures were prepared by 1:100 dilution of overnight cultures into 10 mL of 2X TGY medium (*27*) and incubation at 30 °C with shaking to an OD600 of 0.08-0.16. For colony enumeration, *D. radiodurans* cells were plated on 2X TGY agar plates and incubated at 30 °C until colonies were countable (~48 to 96 hr). Strains used in this study are listed in **Table S3**.

*Serial dilutions and CFU/mL determination*

All serial dilutions were performed in 1X phosphate-buffered saline (PBS) (for 1 L: 8 g NaCl, 0.2 g KCl, 1.44 g $Na_2HPO_4$, $KH_2PO_4$ 0.24 g with 800 mL $dH_2O$, adjust pH with HCl to 7.4, then add remaining 200 mL $dH_2O$**)**. Unless otherwise stated, serial dilutions were performed with serial 1:10 dilutions of 100 µL of culture or previous dilution into 900 µL 1X PBS. Before transfer to the next dilution tube, samples were vortexed for 2 seconds and mixed by pipetting to ensure mixing. One-hundred µL of appropriate dilutions were aliquoted onto agar plates of the appropriate medium and were spread-plated utilizing an ethanol-sterilized, bent glass rod. For spot plating, 10 µL of each dilution was aliquoted onto agar plates of the appropriate medium and spots were allowed to dry before plates were incubated as in *Growth conditions*.

CFU/mL was calculated using the highest CFU count for each strain assayed that remained between 30 and 300 CFU (ex: 250 CFU on a $10^{-4}$ dilution plate would be used for calculation over 40 CFU on a $10^{-5}$ dilution plate).

*Generalized Linac irradiation protocol*

Irradiations were performed as previously described (*41, 42*). Samples were maintained at 4 °C and transported to the University of Wisconsin Medical Rad. Res. Center (UWMRRC) Varian 21EX clinical linear accelerator (Linac) facility for irradiation. The total transport time was approximately 15 min to and from the Linac facility. For each irradiation, the Linac was set to deliver a beam of electrons with 6 MeV of energy to uniformly irradiate all samples (a total of 28) at once. To accomplish this, a special high-dose mode called $HDTSe^-$ was utilized, which resulted in a dose rate to the samples of approximately 72 Gy/min. The sample tubes were placed horizontally and submerged at a depth of 1.3 cm (measured to the center of the tube's volume) in an ice-water filled plastic tank and set to a source-to-surface distance (SSD) of 61.7 cm. A 30 x 30 $cm^2$ square field size was set at the Linac console, which gave an effective field size at this SSD of 18.5 x 18.5 $cm^2$. This is ample coverage to provide a uniform dose to all of the sample vials. The monitor unit calculations (determination of the amount of time to leave the Linac on) were based on the American Association of Physicists in Medicine (AAPM) Task Group 51 protocol for reference dosimetry (*60*). This is the standard method for determining dose per monitor unit in water for radiation therapy calculations. Once the dose was determined in the AAPM Task Group 51 reference protocol conditions (SSD = 100 cm and depth = 10 cm), an ion chamber and water-equivalent plastic slabs were used to translate this dose to the specific conditions used in this project. An independent dose verification was performed with thermoluminescent dosimeters (TLDs) (*41*). TLDs are passive dosimeters that are small, accurate and well-suited for dose verification in the routinely used 1.5 mL sample vials.

*Ionizing radiation resistance assay using the Linac*

Strains were grown in biological triplicate overnight and to an $OD_{600}$ of approximately 0.2 in LB as routinely performed. A 1 mL sample for each dose tested (including 0 Gy) was removed and aliquoted into a sterile 1.5 mL microfuge tube. Samples were pelleted by centrifugation at 13,000 xg for 1 min, and the supernatant was poured off. Samples were resuspended in 1 mL ice-cold 1X phosphate-buffered saline (PBS), and pelleting was repeated. This process was repeated three more times to wash cells. A 100μL aliquot of each culture was removed, serial diluted 1:10 in 900μL of PBS to a final 10,000-fold dilution (with vigorous vortexing of each sample for ~30 sec for undiluted samples, and ~10 sec between each dilution) and 100μL was plated on LB agar to determine the colony forming units (CFU)/mL before irradiation. Samples were maintained at 4 °C and irradiated with the appropriate doses as described. A 100μL aliquot of each culture was removed and plated to determine CFU/mL and percent survival as described.

*Directed evolution protocol using Linac*

Directed evolution was carried out as previously described (*41, 42*). For each round of directed evolution, separate aliquots of 2 mL of LB medium was inoculated with frozen stock of each population from the previous round of selection. These were incubated overnight with aeration at 37 °C and were grown with usual practices in LB medium to an $OD_{600}$ of 0.2 the next day. Each culture was incubated on ice for 10 minutes to stop growth. Two 1 mL samples were removed and aliquoted into sterile 1.5 mL microfuge tubes. Samples were washed three times with 1 mL ice-cold 1X phosphate-buffered saline (PBS) and resuspended in a final volume of 1 mL of 1X PBS. A 100μL aliquot of each culture was removed, serial diluted 1:10 in 900μL of PBS to a final 10,000-fold dilution and 100μL was plated on LB agar to determine the colony forming units (CFU)/mL before irradiation. Samples were maintained at 4 °C and taken to a Varian 21EX clinical linear accelerator (Linac) for irradiation.

After irradiation, an aliquot of each culture was removed, serial diluted 1:10 in 900 μL of PBS to a final 1,000-fold dilution, and 100 μL of each serial dilution was plated on LB agar for each dose to determine the CFU/mL after irradiation. LB agar plates were incubated overnight at 37 °C. Remaining irradiated cultures were pelleted by centrifugation at 13,000xg and supernatant was discarded. These pellets were resuspended in 1 mL of fresh LB medium, and this was added to 1 mL LB medium in a 5 mL glass culture tube. These resuspensions were incubated overnight with aeration at 37 °C. The following day, the percent survival for each dose was calculated using CFU/mL calculations before and after irradiation at each dose. The overnight culture of each population replicate showing closest to 1% survival was stored at -80 °C and used for the next cycle of selection. One cycle of selection was performed weekly due to limited access to the Linac.

The initiating round of selection was done as described above, except the original culture used was an overnight culture of MG1655 prepared from an isolated colony. This protocol was adapted from a previously used protocol (*61*).

*Growth Curves*

Strains were cultured described as in *Growth conditions* overnight and to an $OD_{600}$ of 0.2 in LB medium. Cultures were then diluted 1:10 in 100 μL total of LB medium in a clear, flat bottom 96-well plate (Thermo Fisher Scientific, Waltham, MA; Cat #: 07-200-656**)** and incubated overnight in a Biotek Synergy H1 plate reader **(**Biotek; Winooski, VT**)** at 37 °C with shaking, with $OD_{600}$ readings taken by the plate reader every 10 minutes.

*Confirmation of phenotype stability*

Evolved isolates were serially cultured for 10 days (~100 generations) in the absence of irradiation. Each day (after 24 hr of growth), 30 μL of overnight culture for each isolate was diluted into 3 mL of fresh LB and again incubated overnight at 37 °C. After the final sub-culture grew for 24 hr, 1 mL of culture was mixed with 80 μL DMSO stored at -80 °C as is routinely performed. These frozen stocks were used to generate isolated colonies through streak plating, which were used for irradiation assays to confirm the stability of the radioresistant phenotype.

*Bright-field Microscopy*

One mL of exponential phase culture or 20 μL of overnight culture was washed in 1X PBS as routinely performed; however, the final resuspension volume was 100 μL. Washed cells were maintained at 4 °C until needed. When ready to image, a 10 μL aliquot of each sample was placed on a number 1.5 thickness glass coverslips plate (Thermo Fisher Scientific, Waltham, MA; Cat #: 12-544-EP**)**, and spread by placing a 4% agarose gel pad on the liquid spot. Imaged cells using a Nikon ECLIPSE Ti-E inverted microscope, housed in the University of Wisconsin – Madison Biochemistry Optical Core facility.

*Metals analysis*

Cultured appropriate strains in biological triplicates as described in '*Growth conditions*' to an $OD_{600}$ of 0.2. One mL of each was washed 3 times in 1X Dulbecco's PBS (Sigma-Aldrich, St. Louis, MO; Cat #: D1283) as described in '*Ionizing radiation resistance assay*'. Store-bought PBS was used to avoid the possibility of metals contamination from lab-specific $dH_2O$. One-hundred µL of each sample was diluted and plated as in '*Serial dilutions*' in order to determine the CFU/mL of each sample to normalize relative amounts of quantified elements to cell concentration. The remaining washed samples were transported at 4 °C to the University of Wisconsin State Laboratory of Hygiene Trace Element Research Laboratory for Magnetic Sector inductively-coupled plasma mass spectrometer (ICPMS) analysis.

*Deep sequencing*

Genomic DNA was prepped from overnight cultures prepared from frozen stocks of populations from every even round of selection using the Wizard Genomic DNA Purification Kit (Promega, Madison, WI). Duplicate round 150 DNA samples were submitted to the University of Wisconsin Biotechnology Center for library preparation and whole genome sequencing using a NovaSeq6000 instrument generating paired 150 bp reads. Reads were aligned to the reference genome (U00096.3) and mutations were called using DNAstar SeqMan NGen (Madison, WI) software.

Rounds 1-100 DNA samples were submitted to the Department of Energy Joint Genome Institute (Berkeley, CA) for sequencing and analysis. DNA was randomly sheared into ~500 bp fragments and the resulting fragments were used to create an Illumina library. This library was sequenced on Illumina HiSeq generating 100bp paired end reads. Reads were aligned to the reference genome using BWA (*62*), downsampled to an average depth of 250 fold coverage with picard (http://broadinstitute.github.io/picard) and putative mutations and small indels were called using callvariants.sh from BBMap (sourceforge.net/projects/bbmap). Sequencing results for rounds 1-100 are published (*41, 42*).

Sequencing results for round 150 are reported in their entirety in Supplementary Data 1. However, for analysis of numbers and types of mutations in each population, mutations in genes with high enough homology to have suspected mismapping of reads (i.e. *rrs*, *rrl*, *rrn*, *rhs*, and *ins* genes) were not considered due to increased likelihood of a false-positive mutation call. In addition, mutations with inconsistent frequency calls (ex: jumping from 0, to 100, to 0% allele frequency) were also not used. All mutations removed from consideration are also listed in Supplementary Data 1.

Genomic data from the Joint Genome Institute was obtained as described previously (*41, 42*).

*Label free quantification mass spectrometry with E. coli and D. radiodurans*

Irradiated (3000 Gy) and untreated (0 Gy) samples were lysed by addition of SDS to a final concentration of 2.0%, then immediately subjected to protein extraction and concentration using a standard methanol:chloroform protocol. Purified protein pellets were solubilized in

8M urea with 50mM ammonium bicarbonate (AMBIC) and subjected to a standard BCA assay to determine protein concentration.

For each of the samples, 10 μg (varying volumes) of each was diluted to 4M urea with 50mM AMBIC and treated with 2mM dithiothreitol for 30 min at 50 °C, 5mM iodoacetamide for 30 min at room temperature in darkness, and then 2mM dithiothreitol for 5 min at room temperature. Samples were diluted further to 1M urea with 50mM AMBIC, and 0.05μg of Trypsin and Lys-C proteases were each added (final protease mass:protein mass of 1:100) . Samples were incubated overnight at 37 °C, for 15hr total.

Digestions were stopped with addition of neat formic acid to 1.0%, subjected to solid phase cleanup using Agilent C18 OMIX tips (Agilent Technologies; Santa Clara, CA) according to manufacturer's protocol, and dried down to completion using a vacuum centrifuge.

Samples were injected for analysis using an UltiMate 3000 RSLCnano system (Thermo Fisher; Waltham, MA) onto an Orbitrap Lumos mass spectrometer (Thermo Fisher; Waltham, MA). A 75μm x 50cm Pepmap RSLC column (Thermo Fisher; Waltham, MA) packed with 2μm beads and 100Å pore size was used as the stationary phase. Mobile phase A was 0.1% formic acid and mobile phase B was 80% acetonitrile/0.1% formic acid. A 75-minute elution gradient to 37.5% B was used, after which 95% B was flushed for 5 minutes and column re-equilibration using 2% B was performed for 10 minutes. DDA-MS was performed with the following parameters: MS1 spectra were acquired in profile mode in the Orbitrap with a resolution of 120K and a scan range of 350-1600 m/z. A normalized AGC target of 250% and automatic max inject time was used. Charge state filtering of 2-7, monoisotopic peak selection set to peptide, and dynamic exclusion of 10 seconds, n=1 and with a mass tolerance of +/-10 ppm were used for triggering MS2 acquisition. Cycle time between MS1 scans was set to a max of 1s. For MS2 acquisition, an isolation window of 0.7 Da was used and peptides were fragmented using HCD with a collision energy of 32%. MS2 were acquired in centroid mode in the Ion Trap using the automatic scan range parameter and scan rate set to turbo. An AGC target of 3e4 and an automatic max inject time were used.

Data was analyzed using the Sequest algorithm within Proteome Discoverer (PD) (Thermo Fisher; Waltham, MA). For *E. coli* searches, the Uniprot K12 *E. coli* proteome, downloaded on 7/2/2019, was used (PID: UP000000625, 4382 sequences including contaminants). For *D. radiodurans* searches, the Uniprot *D. radiodurans* proteome, downloaded on 3/27/2018, was used (PID: UP000002524, 3172 sequences including contaminants). Databases were searched with the specified parameters: trypsin with 2 possible missed cleavages, precursor and fragment mass tolerance 10 ppm and 0.6 Da, respectively, and a max amount of 4 dynamic modifications per peptide. Dynamic modifications were specified as carbamidomethyl/+57.021Da (on C), oxidation/+15.995Da (on CDEFHILMNPQRSTVWY), carbonylation/+13.979Da (on ACDEFHILKMNPQRSTVWY), dioxidation/+31.990Da (on ACDEFHIKLMNPRSTVWY), and trioxidation/+47.985 Da (on CFWY). No static modifications were set. Searches were based on previous reports of abundance of the given modifications on each amino acid reside (*12*). A false discovery rate (FDR) for peptide spectral matches (PSMs), peptides, and proteins of 0.05% was used via percolator in PD. For quantification, a combination of the Minora Feature Detector, Feature Mapper, and Precursor Ions Quantifier nodes were used in PD. Samples were grouped per strain or organism as 5 untreated replicates untreated vs. 5 treated replicates. Default settings were used for the Minora Feature Detector and Feature Mapper nodes. For the Precursor Ion Quantifier node, Intensity was used for precursor quantification and normalization was performed using total peptide amount per file. Both unique and razor peptides were used to quantify protein level differences, excluding modified peptides. Protein levels were quantified using summed abundances, no data imputation, and ANOVA hypothesis testing on individual proteins. The Benjamini-Hochberg method was used to adjust p values, and the adjusted p values which were used for significance thresholds.

#### Supplemental Figures

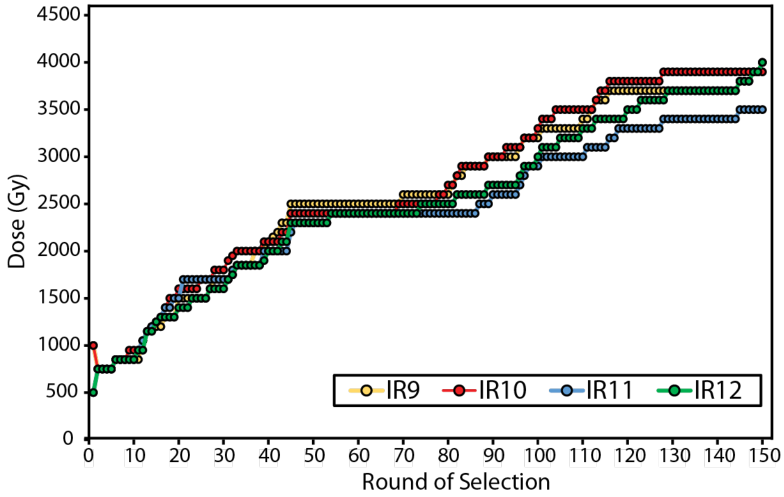


**Figure S1. Experimentally evolved ionizing radiation resistance has continued to increase over 150 cycles of selection.** Dose required to kill 99% of each population has increased. Each data point indicates the dose of IR that each population was given prior to being outgrown overnight and stored at -80° C. The percent survival was estimated from a single replicate at each round of selection.

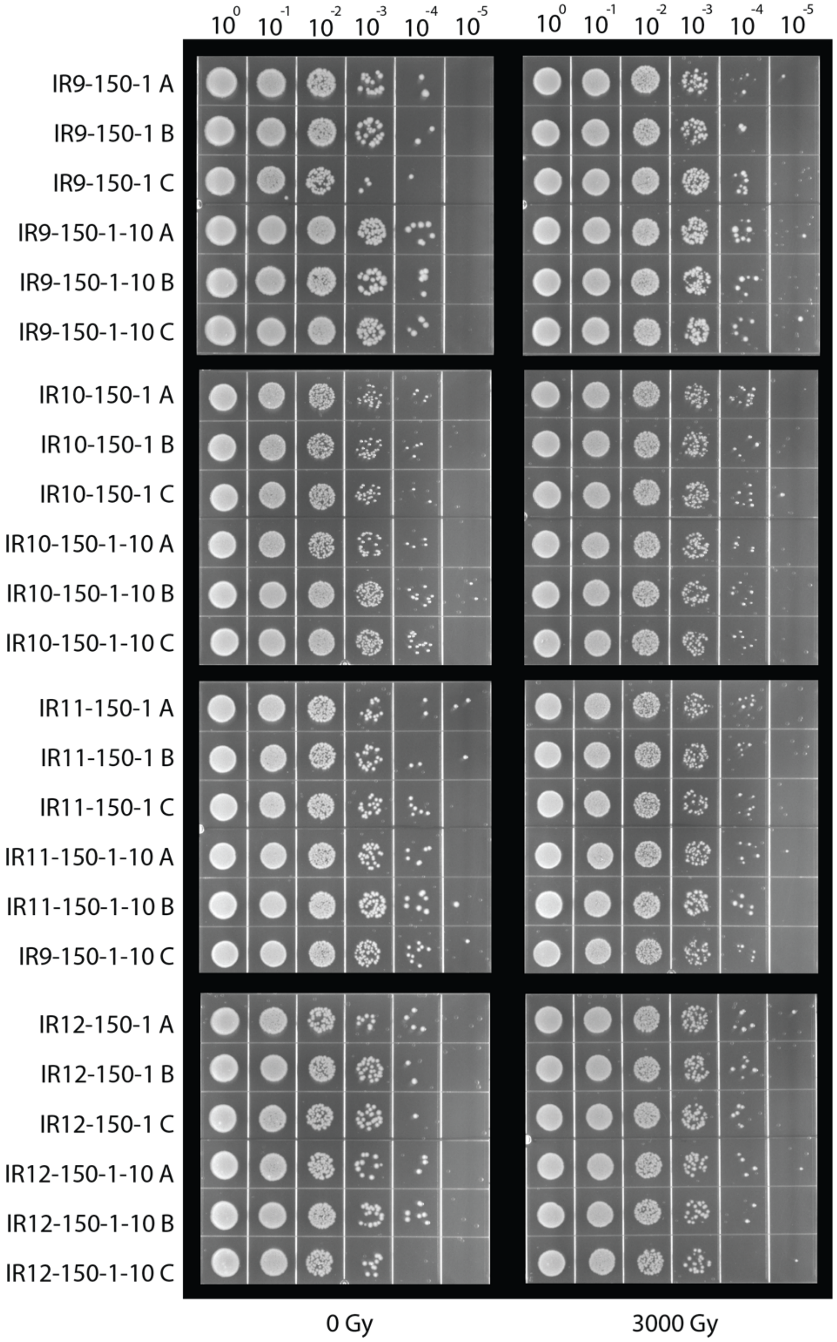


**Figure S2. The radioresistant phenotype of the evolved isolates is stable in the absence of selection.** Isolates were serial-cultured for 10 days (approximately 100 generations) as described in the *Materials and methods*. Isolated colonies from streak-plated freezer stocks of these newly-generated isolates (named IRX-150-X-10; ex: IR9-150-1-10 is IR9-150-1 after 10 days of serial culture without selection) were used for the irradiation assays. No difference in IR resistance was observed. The data are representative of biological triplicates for each strain from two independent experiments.

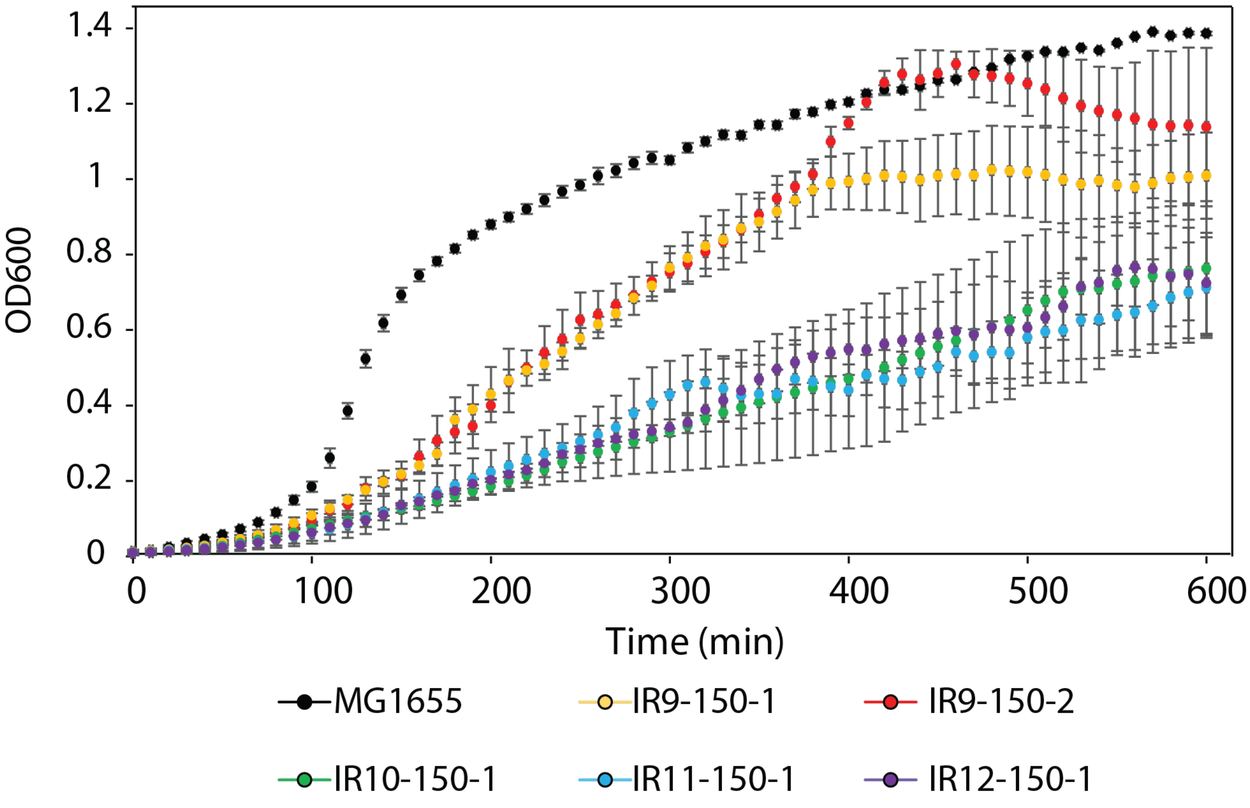


**Figure S3. Growth curves of evolved isolates reveal significant growth defects.** MG1655 is the Founder strain used to begin the evolution experiment. Strains named IR'X'-150-1 are isolates from the indicated mixed population at 150 cycles of selection. IR9-150-2 is a second isolate from population IR9-150, and represents a sub-population separate from IR9-150-1. Cultures of indicated strains were grown in LB medium overnight and then to early exponential phase as described in the *Materials and methods*. All cultures were incubated at 37 °C during growth. Early exponential phase cultures were diluted 1:10 in a final volume of 100 µL of LB growth medium, and then incubated overnight in a Biotek Synergy H1 plate reader, with $OD_{600}$ measurements taken automatically every 10 minutes. This experiment is representative of two independent experiments performed in biological triplicate; error bars represent the standard deviation of the $OD_{600}$ measurements of a single biological triplicate.

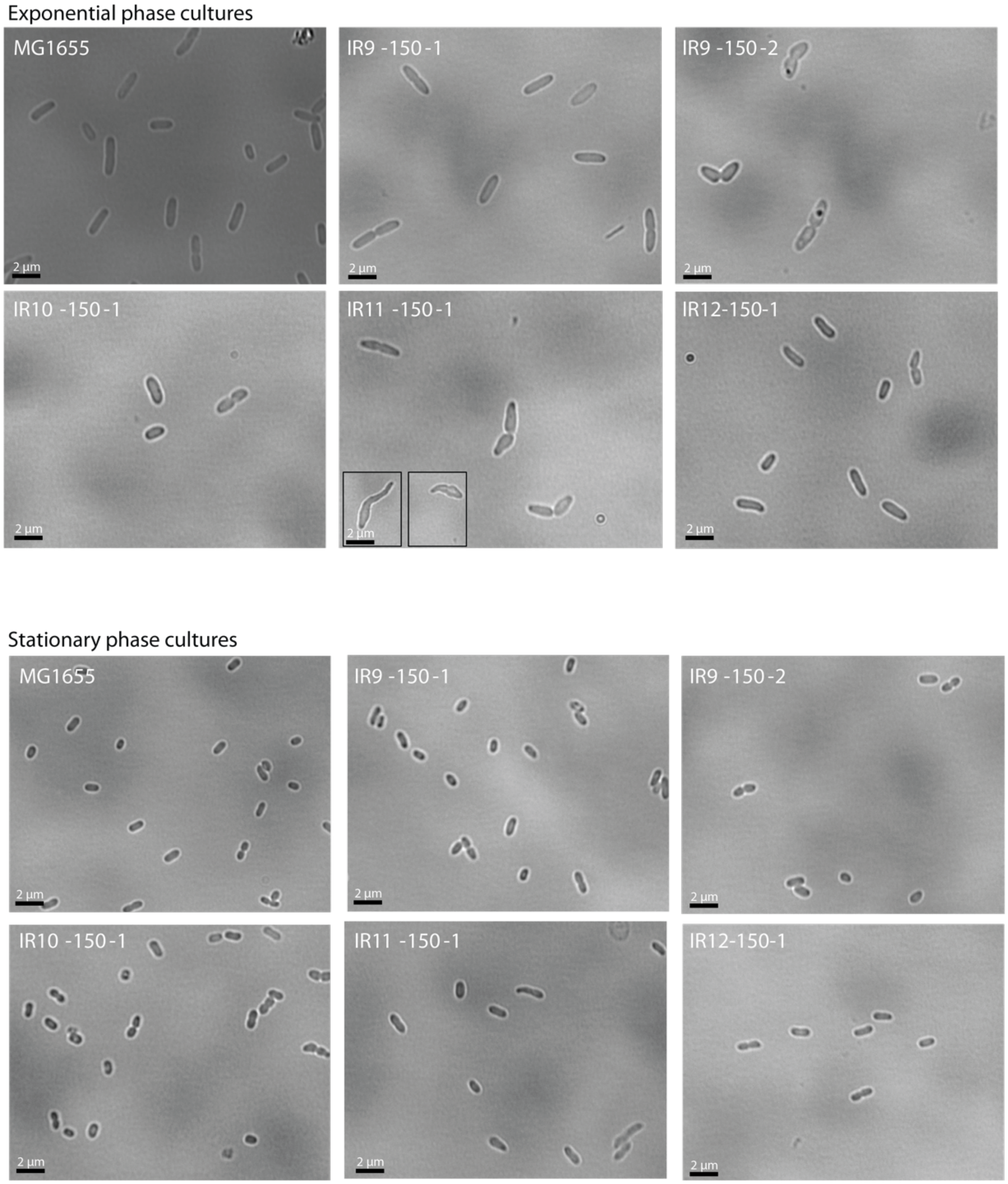


**Figure S4. Microscopy reveals significant changes to cell morphology of evolved isolates**. Exponential and stationary phase cultures were imaged as described in the *Materials and methods*. The 2 µm scale bar was determined using the length of a non-dividing, exponential phase MG1655 cell and applied to all images (which were taken at 1000x magnification using immersion oil). The dark circles inside cells are unique to IR9-150-2; the nature of these structures is not yet known. The morphologies of IR11-150-1 cells are particularly divergent from MG1655. Two inserts from different images of IR11-150-1 were included to demonstrate the variation in IR11-150-1 morphology. Images are representative of two independent experiments, where greater than 200 cells were observed for each isolate.

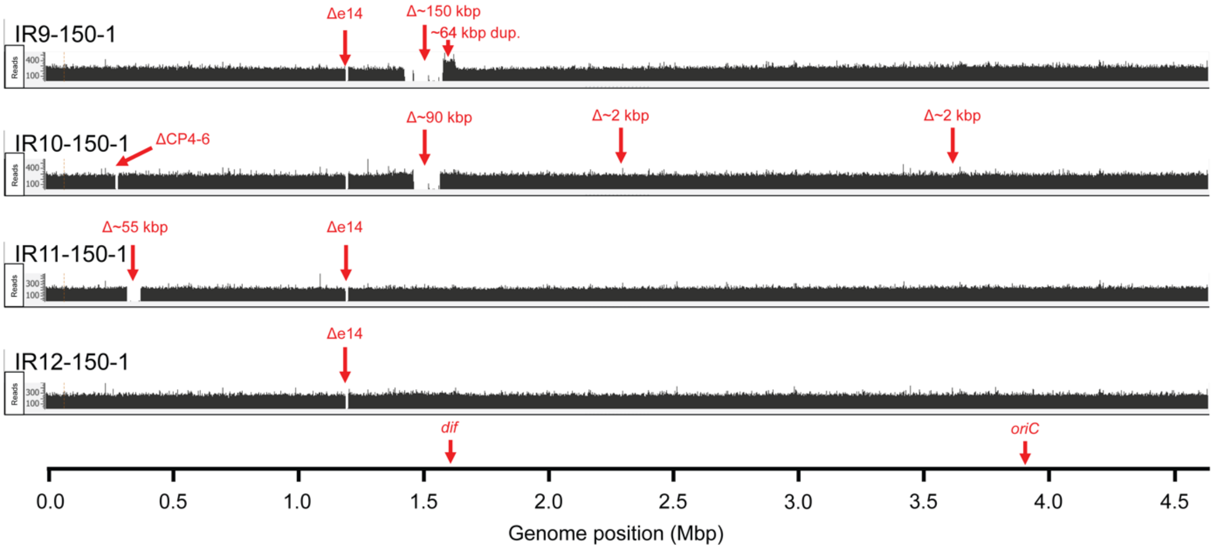


**Figure S5. Major deletions and duplications in evolved populations.** The ~55 kbp deletion in IR11 is new. The region deleted includes the *lac* operon. Deletions and duplications shown for IR9 and IR10 were reported previously (42).

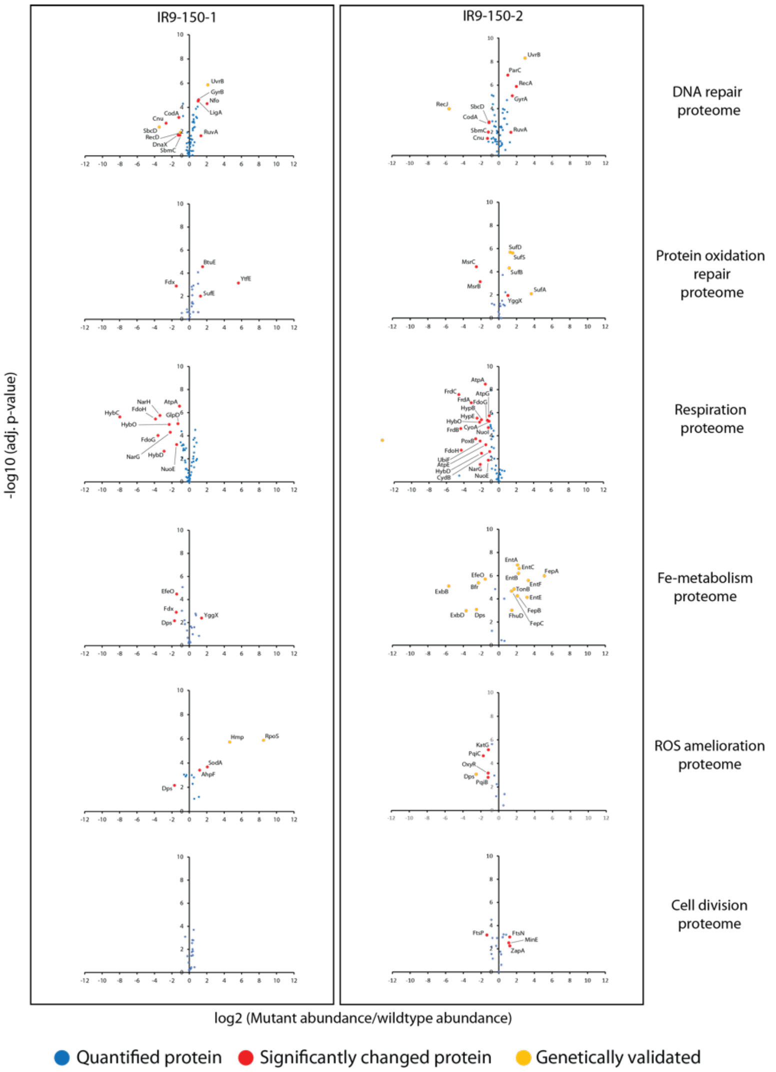


**Figure S6. The proteome composition of IR9-150-1 and IR9-150-2 have diverged from *E. coli* MG1655. Data broken down by function.** Volcano plots depict each protein detected with a circle, with the fold difference in abundance compared to the same protein in MG1655 on the x-axis and the p-value of that change on the y-axis. Proteins with a significant fold change are colored in red. The circle is colored in orange if a mutation in the appropriate evolved isolate could reasonably explain the change in protein level (i.e. a frameshift in gene X may lead to lowered abundance of protein X, or increased abundance of protein Y if protein X regulates levels of protein Y). The proteome composition of MG1655, IR9-150-1, and IR9-150-2 was determined utilizing label free quantification mass spectrometry (LFQ-MS). Significant changes in protein abundance were defined as an increase or decrease greater than two-fold, with adjusted p-values less than 0.05 (calculated using Benjamini-Hochberg correction). Only proteins with more than one detected peptide were considered for our analyses. Supplemental Datasets 1 and 2 contain the genomics and proteomics data used for these analyses, respectively.

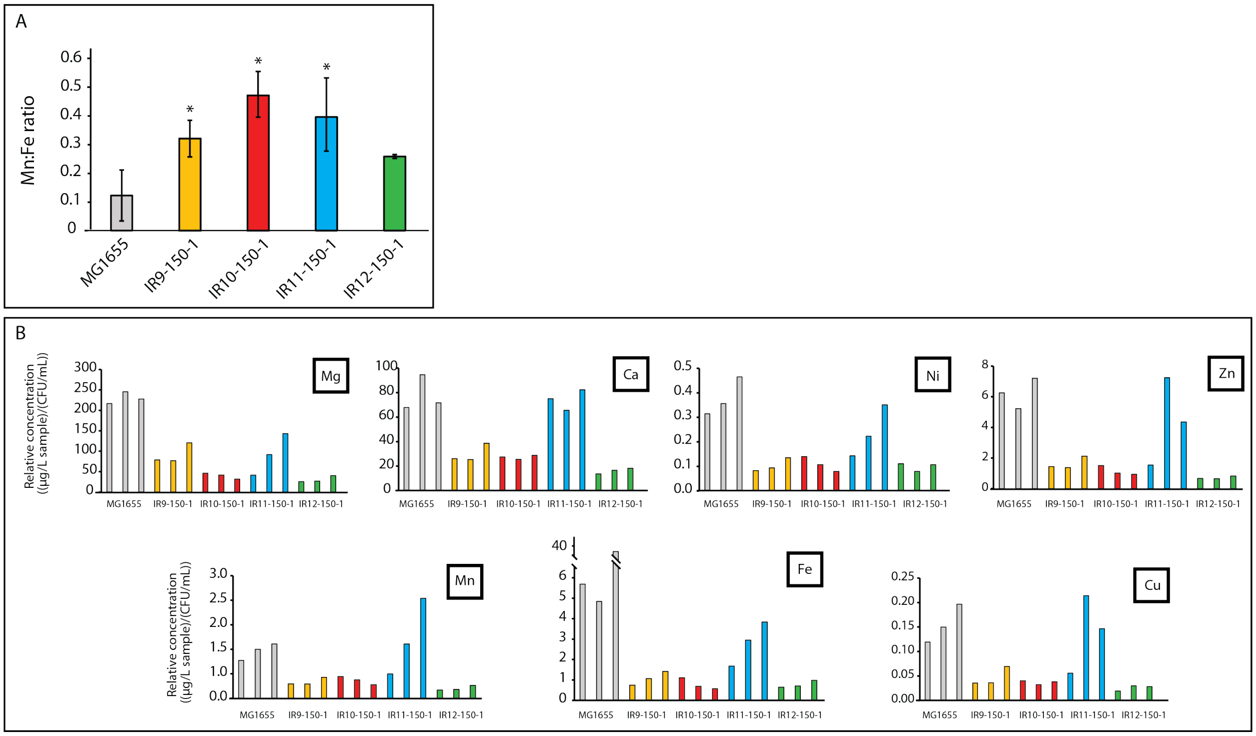


**Figure S7. Metals analysis of evolved *E. coli* isolates after 150 cycles of selection**. A) The Mn:Fe ratio was calculated for each isolate tested. Small but significant ( p-value > 0.05, indicated by the '*' symbol) differences between the Mn:Fe ratio of the evolved strain and MG1655 were observed. B) Concentrations of trace metals were analyzed in early exponential phase cultures of the noted strains by the University of Wisconsin State Hygiene Laboratory of Hygiene Trace Element Research Laboratory. Raw concentrations of trace elements for each biological replicate for each strain was normalized to the CFU/mL of each culture. These data represent the results of biological triplicates for each strain listed.

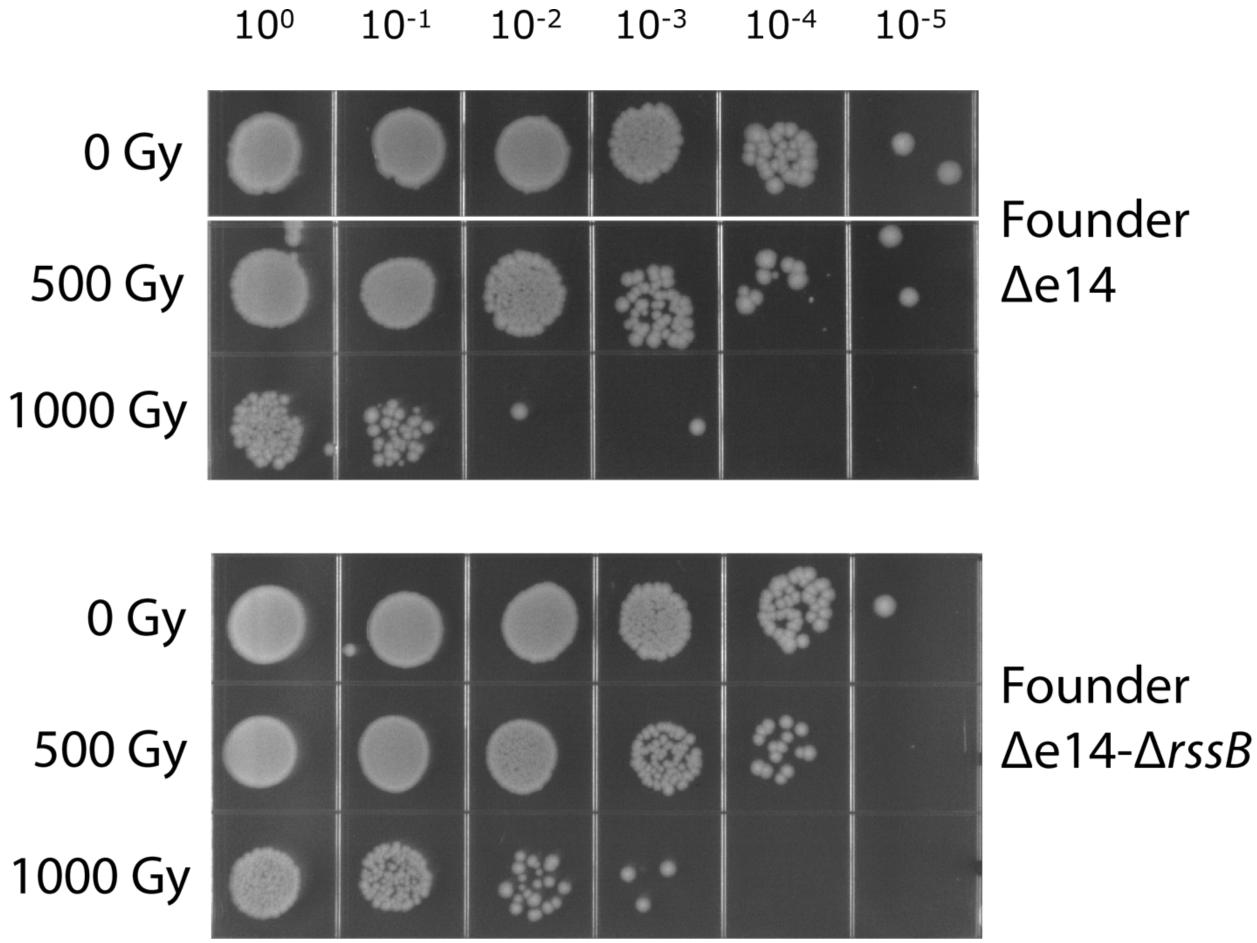


**Figure S8. Contribution of an *rssB* deletion to IR resistance in a wild type background.**

#### Supplemental Tables

| | Parent bp | Mutant bp | IR9-150-1 | IR9-150-2 | IR10-150-1 | IR10-150-2 | IR11-150-1 | IR11-150-2 | IR12-150-1 | IR12-150-2 |
|---|---|---|---|---|---|---|---|---|---|---|
| Transitions | AT | GC | 187 | 225 | 156 | 175 | 121 | 130 | 140 | 127 |
| | GC | AT | 296 | 351 | 264 | 210 | 275 | 250 | 339 | 330 |
| Transversions | AT | CG | 48 | 49 | 28 | 41 | 31 | 29 | 37 | 40 |
| | GC | TA | 214 | 240 | 164 | 159 | 148 | 136 | 235 | 227 |
| | AT | TA | 130 | 154 | 73 | 75 | 74 | 73 | 104 | 104 |
| | GC | CG | 57 | 67 | 49 | 44 | 45 | 54 | 60 | 58 |
| Totals | Transitions | | 483 | 576 | 420 | 385 | 396 | 380 | 479 | 457 |
| | Transversions | | 449 | 510 | 314 | 319 | 298 | 292 | 436 | 429 |
| | Ts:Tv | | 1.08 | 1.13 | 1.34 | 1.21 | 1.33 | 1.30 | 1.10 | 1.07 |
| Coding | Synonymous | | 236 | 259 | 193 | 190 | 179 | 179 | 230 | 210 |
| | Non-synonymous | | 652 | 776 | 492 | 473 | 451 | 429 | 618 | 610 |
| | Stop gained | | 30 | 33 | 14 | 27 | 20 | 20 | 25 | 23 |
| | Stop lost | | 2 | 3 | 1 | 1 | 3 | 1 | 2 | 2 |
| | Start lost | | 0 | 9 | 4 | 2 | 0 | 0 | 3 | 4 |
| | Insertions | +1 | 2 | 2 | 3 | 2 | 3 | 2 | 5 | 4 |
| | | +2 | 1 | 0 | 0 | 1 | 0 | 0 | 1 | 1 |
| | Deletions | -1 | 89 | 105 | 45 | 58 | 30 | 29 | 61 | 59 |
| | | -2 | 4 | 3 | 3 | 1 | 1 | 3 | 1 | 2 |
| | | -3 | 1 | 1 | 0 | 0 | 0 | 0 | 0 | 1 |
| | | -5 | 0 | 0 | 0 | 0 | 0 | 0 | 0 | 0 |
| | | -9 | 0 | 0 | 0 | 0 | 0 | 0 | 0 | 0 |
| | Total | | 1017 | 1191 | 755 | 755 | 687 | 663 | 946 | 916 |
| Non-coding | SNPs | | 169 | 190 | 117 | 132 | 112 | 108 | 170 | 166 |
| | Insertions | +1 | 1 | 1 | 2 | 0 | 0 | 1 | 0 | 1 |
| | Deletions | -1 | 17 | 18 | 10 | 19 | 9 | 6 | 16 | 15 |
| | | -2 | 0 | 0 | 0 | 0 | 0 | 0 | 2 | 3 |
| | Total | | 187 | 209 | 129 | 151 | 121 | 115 | 188 | 185 |
| **Overall Total** | | | **1204** | **1400** | **884** | **906** | **808** | **778** | **1134** | **1101** |

**Table S1. Number of polymorphisms in sequenced isolates from each population at round 150.**

| Protein | Description | IR9-150-1 | IR9-150-2 | IR10-150-1 | IR10-150-2 | IR11-150-1 | IR11-150-2 | IR12-150-1 | IR12-150-2 |
|---|---|---|---|---|---|---|---|---|---|

| | | | | | | | | | |
|---|---|---|---|---|---|---|---|---|---|
| ArcB | sensor histidine kinase involved in regulation of anaerobic metabolism | I4N | V38fs | D166Y | - | S74W | | S24P | |
| | | - | - | - | - | R100H | | - | - |
| BluF | blue light- and temperature-regulated antirepressor | V268L | - | - | - | E368A | | R186W | |
| | | | | | | - | - | E261K | |
| | | | | | | S256F | - | S318P | |
| Cas3 | CRISPR-associated endonuclease/helicase | E820G | I561T | Q292. | - | - | - | D848N | |
| | | - | - | P512L | - | - | - | - | E843K |
| CopA | soluble Cu+ chaperone | V270F | | T84A | - | T525A | | A812V | |
| | | - | P198T | - | - | - | - | - | - |
| | | - | H409Q | - | - | - | - | - | - |
| CstA | carbon starvation protein A | A240T | M353K | R73fs | - | - | L204P | - | - |
| | | - | - | R198I | - | - | - | - | - |
| CusA | copper/silver export system | G757C | - | - | - | P371fs | E731A | - | - |
| | | F862L | - | - | - | E445. | - | - | - |
| CvrA | putative K+:H+ antiporter | V394fs | - | V569I | L461M | - | - | E415. | |
| | | - | - | - | - | - | - | R523L | - |
| Dcp | peptidyl-dipeptidase | - | - | A78fs | R302L | A229S | - | A194S | |
| | | - | - | - | M671I | - | - | - | - |
| EntF | apo-serine activating enzyme in enterobactin synthesis | - | A55V | W652. | - | - | T159I | L555P | |
| | | - | A302T | V1083E | - | - | - | P588L | |
| EvgS | sensor histidine kinase involved in acid tolerance | - | A717E | - | - | - | G211C | G491E | |
| | | - | - | - | - | - | A863T | K629R | |
| ExoX | exonuclease X | .221Y | - | S44fs | W96C | G15R | T54fs | A79E | |
| | | - | - | I150fs | - | - | P207Q | A162V | |
| | | - | - | - | - | - | - | V184A | |
| FhuE | ferric coprogen outer membrane transport complex | - | V82fs | N359Y | | T137I | | E334K | |
| | | - | - | D133N | S81L | A297V | - | - | - |
| | | - | - | K242Q | - | - | - | - | - |
| FimE | DNA recombinase, regulator for fimA | I125F | - | R154P | R178C | - | G181V | R59fs | |

| | | | | | | | | | |
|---|---|---|---|---|---|---|---|---|---|
| | | H136Y | - | - | - | - | - | E184fs | |
| FlgK | flagellar hook-filament junction protein 1 | N368Y | S83G | E514K | - | M229T | - | - | - |
| | | Q523H | - | - | - | - | - | - | - |
| FtsK | cell division DNA translocase | - | - | P961S | - | T54M | P787L | D724N | |
| | | - | - | - | - | - | - | Q802R | |
| GsiB | glutathione ABC transporter | - | A120V | - | N345fs | D197E | | F240fs | |
| | | - | A509G | - | - | - | - | A241G | |
| Gss | fused glutathionylspermidine amidase / synthetase | - | **E608Q** | V94A | Y274fs | V552A | | - | - |
| | | - | - | **E608Q** | - | A63S | - | T3N | |
| Kup | K+:H+ symporter | A78D | - | - | L585F | - | - | E229D | |
| | | - | - | - | - | - | - | - | S334P |
| Lhr | putative ATP-dependent helicase | - | T1505M | H710D | - | G1534V | W1236. | A716G | |
| | | - | - | A808T | - | - | - | - | - |
| | | - | - | S1256P | - | - | - | - | - |
| | | - | - | R1400H | - | - | - | - | - |
| LldP | lactate/glycolate:H+ symporter | - | G329V | F369L | - | - | A330S | - | S462. |
| | | - | - | - | - | - | - | - | Q477. |
| MdtC | multidrug efflux pump permease subunit | - | F898L | - | N936D | - | - | R99Q | |
| | | - | - | - | - | - | - | P325S | |
| | | - | - | - | - | - | - | P754L | |
| MdtG | multidrug efflux pump | L120H | | S354P | - | - | - | - | - |
| | | - | M54I | - | - | - | - | - | - |
| | | - | T146fs | - | - | - | - | - | - |
| | | - | A234fs | - | - | - | - | - | - |
| MdtK | multidrug efflux pump | A57E | | P265fs | F60L | V351I | - | - | - |
| | | - | - | - | R337W | - | - | - | - |
| MscM | miniconductance mechanosensitive channel | Q128K | | A105G | - | R287C | - | - | - |
| | | - | H364Q | - | - | - | - | - | - |
| | | - | G689fs | - | - | - | - | - | - |

| | | | | | | | | | |
|---|---|---|---|---|---|---|---|---|---|
| NrdD | anaerobic nucleoside-triphosphate reductase activator | - | R525S | - | Y573C | G709E | | - | - |
| | | - | - | - | - | - | R455fs | - | - |
| | | - | - | - | - | - | Y573C | - | - |
| | | - | - | - | - | - | F605S | - | - |
| Nth | endonuclease III | V57M | - | V36fs | - | T43fs | | K85N | |
| | | - | - | I79L | - | - | - | - | - |
| | | - | - | C203Y | - | - | - | - | - |
| PflD | putative formate acetyltransferase 2 | I583F | - | T16I | - | E602fs | - | - | - |
| | | D584E | - | L408M | - | - | - | - | - |
| PgaA | β-1,6-N-acetyl-D-glucosamine outer membrane porin | Q309. | | E531. | S321L | - | P28A | L257P | |
| | | - | - | - | - | - | - | - | E484D |
| PqiA | SoxRS-regulated intermembrane transport protein | A367T | | - | - | - | - | A21V | - |
| | | - | P176S | - | - | - | - | W177L | - |
| | | - | - | - | - | - | - | W197. | - |
| ProB | glutamate 5-kinase | N116fs | | S308fs | - | - | - | R291L | |
| | | - | A200T | - | - | - | - | - | - |
| | | - | R314C | - | - | - | - | - | - |
| ProX | glycine betaine ABC transporter | W86. | P203Q | Y71C | - | N178S | - | - | - |
| | | D120G | - | - | - | - | - | - | - |
| | | Y127C | - | - | - | - | - | - | - |
| PtsP | phosphoenolpyruvate-protein phosphotransgerase | D28G | - | R119fs | | - | L58M | D641G | |
| | | - | - | - | - | - | R526C | - | - |
| PuuP | putrescine:H+ symporter | I94F | - | V77I | - | - | - | - | - |
| | | N95fs | - | V329A | - | - | - | - | - |
| | | - | - | A354P | - | - | - | - | - |
| RapA | RNA polymerase-binding ATPase and RNAP recycling factor | R131P | R87C | A959V | W779. | - | - | V856fs | |
| | | - | E940K | - | D958G | - | - | - | - |
| RcsD | RcsD-N-phospho-L-histidine | D568N | - | Y212F | H332Q | N66K | | **I380L** | - |

| | | | | | | | | | |
|---|---|---|---|---|---|---|---|---|---|
| | | - | - | G588fs | **I380L** | - | - | - | - |
| RecD | exodeoxyribonuclease V | A90E | | N124D | | A550E | | S92I | |
| | | - | G362D | Q463. | | - | C442R | - | - |
| | | - | - | - | Y418D | - | - | - | - |
| RecJ | ssDNA-specific exonuclease | .578L | | G502D | - | A432S | D166N | F426L | |
| | | M360I | - | - | - | - | - | - | - |
| RecN | SMC-like DNA DSB repair protein | K429Q | | - | R415L | R102P | | S318R | |
| | | - | - | - | - | - | - | A361T | |
| RhlB | ATP-dependent RNA helicase | K27R | | - | W255fs | - | - | F302C | |
| | | - | E254K | - | - | - | - | - | Q160fs |
| RibD | fused deaminase/reductase involved in riboflavin biosynthesis | S193I | Y6H | - | - | - | G119D | I353V | |
| | | - | W297R | - | - | - | - | P264Q | - |
| RpoB | RNA polymerase beta subunit | S72N | | V630E | E244V | P535L | | T600I | |
| | | - | - | K1200E | A977D | S574F | | - | P1181S |
| SbcC | ATP-dependent structure-specific DNA nuclease | - | D661fs | R146S | | Q98K | Q617. | Q861. | R102fs |
| | | - | - | - | F147I | - | L902M | - | - |
| ThrS | threonine—tRNA ligase | P188S | | E326K | | - | - | - | - |
| | | E258D | - | A32V | - | - | - | - | - |
| | | I514S | - | D49E | - | - | - | - | - |
| UvrD | DNA helicase II | I572S | - | A599V | E117K | .721Q | A196D | F702Y | - |
| YbaT | putative transporter | - | S253. | R111C | T289M | A362P | - | - | - |
| | | - | D262N | I163F | - | - | - | - | - |
| YbhJ | putative hydratase | R597fs | | - | L479S | - | - | - | L54I |
| | | C649Y | L576fs | - | - | - | - | - | - |
| YdjE | putative transporter | R117P | | - | G121fs | - | S415. | - | - |
| | | E199G | - | - | T122A | - | - | - | - |
| YeaG | protein kinase | - | - | P398L | F4S | S336F | - | R32fs | |
| | | - | - | - | - | P572T | - | - | - |
| YeeJ | inverse autotransporter adhesin | - | - | - | R866W | D255V | | - | D1651fs |

| | | | | | | | | | |
|---|---|---|---|---|---|---|---|---|---|
| | | - | - | - | S2324C | A713S | | - | - |
| YfaL | putative autotransporter adhesin | R630fs | | - | Q882E | P1177R | | A675T | |
| | | L770F | | - | N883fs | A631V | - | - | - |
| | | G326V | - | - | - | - | - | - | - |
| YfbS | putative transporter | - | - | V8A | R215. | - | - | V185E | |
| | | - | - | - | - | - | - | K359N | |
| | | - | - | - | - | - | - | S567P | |
| YhbX | putative hydrolase | W15. | G34R | - | - | A35E | | - | - |
| | | - | L61Q | - | - | L344V | - | - | - |
| YpjA | adhesin-like autotransporter | - | - | M1388I | - | - | G71fs | T116I | |
| | | - | - | - | - | - | T101A | E200K | |
| | | - | - | - | - | - | - | G910R | - |

**Table S2. Commonly mutated genes across all sequenced round 150 isolates reveal potential drivers of IR resistance.** Genes listed are those with at least 5 unique non-synonymous mutations present across all 8 sequenced isolates.

| Strain | Relevant Genotype | Source |
|---|---|---|
| MG1655 | YbhJ L54I + MntP G25D + RIP321 A 4296380* ACG + GlpR C 3560455* CG + GatC ACC 2173360* A | Blattner, 1997 |
| EAW7704 (Founder Δe14) | MG1655 Δe14 + RbsR L92R + CytR Q110 stop + *fabI/ycjD* int (G 1351174* A) + *yifN/ppiC* int (T 3959934* C) | Harris, 2009 |
| IR9-150 | MG1655 exposed to 150 iterative rounds of IR; mixed population | This study |
| IR10-150 | MG1655 exposed to 150 iterative rounds of IR; mixed population | This study |
| IR11-150 | MG1655 exposed to 150 iterative rounds of IR; mixed population | This study |
| IR12-150 | MG1655 exposed to 150 iterative rounds of IR; mixed population | This study |
| IR9-150-1 | Isolate from IR9-150 | This study |
| IR9-150-2 | Isolate from IR9-150 | This study |
| IR10-150-1 | Isolate from IR10-150 | This study |
| IR10-150-2 | Isolate from IR10-150 | This study |
| IR11-150-1 | Isolate from IR11-150 | This study |
| IR11-150-2 | Isolate from IR11-150 | This study |
| IR12-150-1 | Isolate from IR12-150 | This study |
| IR12-150-2 | Isolate from IR12-150 | This study |
| EAW1480 | IR9-150-1 + RssB Q3 wt | This study |
| STB264 | IR9-150-2 Δ*rssB* | This study |
| STB265 | IR10-150-1 Δ*rssB* | This study |
| *Deinococcus radiodurans* R1 | N/A | Anderson, 1956 |
| * nucleotide position compared to NCBI GenBank U00096.3 reference sequence | | |

**Table S3. Strains used in this study** (*41, 42, 57, 61*).

#### Supplementary Datasets

**Supplementary Dataset 1.** Complete listing of mutations detected in the round 150 evolved isolates.

**Supplementary Dataset 2.** Proteome composition of MG1655 compared to IR9-150-1 and IR9-150-2.
